# A Hidden Markov Model to Estimate the Time Dairy Cows Spend in Feeder Based on Indoor Positioning Data

**DOI:** 10.1101/250092

**Authors:** Matti Pastell, Frondelius Lilli

**Affiliations:** Natural Resources Institute Finland (Luke), Production Systems, Koetilantie 5, 00790 Helsinki, Finland

**Keywords:** Indoor positioning, Feeding time, hidden Markov model, Ultra wide-band, Dairy cow

## Abstract

The feeding time of dairy cows is linked with the health status of the animal and can be used to estimate daily feed intake together with other measurements. The aim of this study was to develop a model to measure the time a dairy cow spends at a feed bunk using an Ultra wide-band indoor positioning system.

We measured the feeding behavior of 50 dairy cows during 7 days using Ubisense indoor positioning system and Insentec roughage feeders. We calculated the feeding (presence at the feeder) probability of the cow using logistic regression model with the distance to feed barrier as input and used the Viterbi algorithm to calculate the most likely state (feeding or not feeding) given state transition probabilities. The model was able to predict whether the cow was at the feeding trough or not with the accuracy of 97.6%, sensitivity 95.3% and specificity 97.9%. The model was also able to estimate the mean bout duration and the number of feeding bouts.

## 1 Introduction

The feeding time of dairy cows is a good indicator of animal status. The time that the cows spend at the feeding barrier has been shown to be linked with lameness (Norring et al., 2014), metritis (Urton et al., 2005) and has been used to improve the estimation of cows’ daily dry matter intake (Halachmi et al., 2016).

Several indoor positioning systems have been introduced for use on commercial dairy farms. Ultra wide-band (UWB) based systems have an accuracy of below 1m in dairy barns after proper filtering (Pastell et al., 2018; Porto et al., 2014). These systems have been used to measure the feeding time of cows based on their proximity to feeding area (Shane et al., 2016; Tullo et al., 2016; Oberschätzl et al., 2015) as compared to behavioral observations. Other automatic on-farm options for monitoring feeding time include RFID based systems, accelerometers (Arcidiacono et al., 2017; Thorup et al., 2016) and computer vision (Porto et al., 2015).

The aim of this study was to develop a model to accurately measure the time at the feed bunk, visit duration and number of visits using an indoor positioning system data. A hidden Markov model was developed and the model performance was compared against reference data from automatic feed intake measurement system.

## 2 Materials and Methods

### 2.1 Data collection

#### 2.1.1 Measuring feeding time

The time that the cows spend at feeding troughs was measured using Insentec (Hokofarm Group B.V., Marknesse, The Netherlands) roughage intake control (RIC) system. The system has a feeding trough where the feed is delivered, a photocell to detect that the cow’s head is located in the trough and an RFID reader to identify the animal (Figure 1). It records the start and end time of each visit and the weight of the consumed feed. The feeders were equipped with barriers to prevent stealing behavior (Ruuska et al., 2014).

**Fig. 1.**
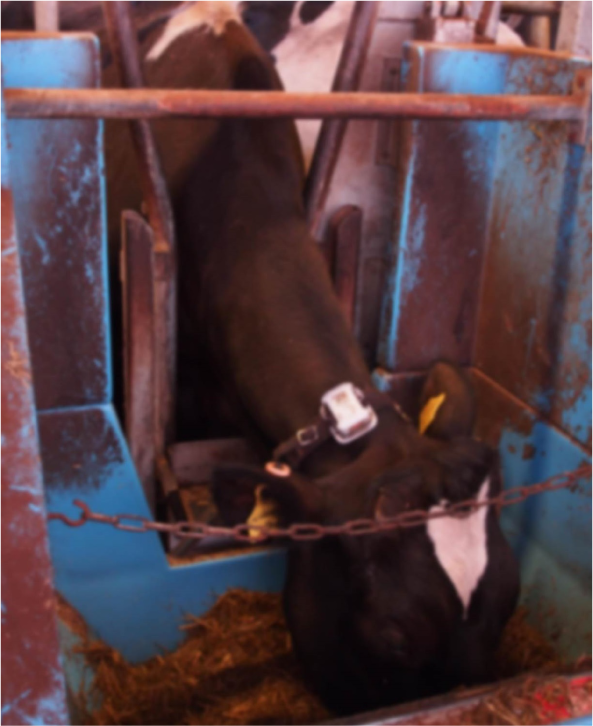
Cow visiting Insentec RIC feeder trough with Ubisense positioning tag attached to the neck collar.

#### 2.1.2 Indoor positioning data

We followed 50 dairy cows of Nordic red and Holstein breeds in a freestall barn using using Ubisense (Ubisense GmbH, Düsseldorf, Germany) UWB-based indoor positioning system. The system recorded the position of each cow at 1.2Hz. Positioning tags (Ubisense Series 7000 Industrial tag) were mounted on cow neck collars (Figure 1) with the tag is positioned on the top of the neck. The tags transmitted UWB pulses to remote sensors mounted on the wall. The Ubisense system calculated the location of the tags using time-difference of arrival technique. The positioning system was set up to cover an area of 22 × 25 m with six sensors and calibrated for the area according to the system manual. The roof height of the barn was between 2.8 m and 7.5m.

The data was recorded using custom measurement software written in Java that stored the data in HDF5 format in a separate file for each day. Raw Ubisense data contained unwanted noise and missing data and was filtered using a heuristic jump filter developed for the system combined with a 5th order median filter. The missing data was interpolated using a piecewise constant interpolation. After filtering the positioning error of the system was below 1m except for the corners of positioning area where the worst case error was up to 2m. The height measurement (z-axis) was not calibrated and therefore not used in the analysis. The system setup, and validation and the filtering method are described in detail in (Pastell et al., 2018).

Cows were housed in two sections of 24 cows in a freestall curtain-wall barn with rubber mattresses and steel separators in stall and, slatted alleys cleaned using manure robots. Both sections had their own concentrate feeder and 12 Insentec RIC-feeders. The cows had free access to total mixed ratio, including grass silage with concentrate (barley-rapeseed meal mixture 80:20) achieving 10 MJ/kg in dry matter. Feed was delivered five times a day using an automatic feeding robot and cows milked on a herringbone milking parlour 2 times a day.

### 2.2 Hidden Markov Model for Classifying Feeding

The collected positioning data was split into training and validation sets, where training set consisted of data from 25 cows and the validation dataset from different 25 cows collected during 7 measurement days. The model was fitted using the training set and the model performance was evaluated using the validation set. The training dataset consisted of 16.9 million observations and the validation dataset of 15.9 million observations.

We used a Hidden Markov model (HMM) to calculate whether a cow is feeding or not feeding for each observation. HMMs can be used to estimate the unknown state of a process based on observed measurements. The model needs a probability for each state given the observations (emission probability) and state transition probability which encodes the probability of the process changing from one state to another (Zucchini et al., 2016).

In this case the model has only two states and the state vector *θ* was:

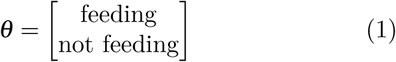

The state transition matrix was obtained directly from Insentec feeder data from teaching dataset:

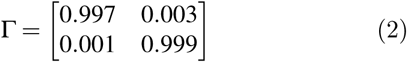

In order to obtain emission probabilities (probability of feeding given cow position) we calculated the Euclidean distance to the feed barrier d_b_ for each positioning sample. Samples on the side of the feed were defined as negative and on the side of the cows positive. We fitted a logistic regression model using the distance to feedbunk as predictor variable in order to calculate the probability of feeding for each sample.

The most likely state was then calculated using Γ and emission probabilities from the fitted logistic regression model using the Viterbi algorithm (Viterbi, 1967). The Viterbi algorithm is a recursive method to calculate the most likely path of hidden states given observed probabilities and state transition probabilities.

Data was analyzed using R 3.3.1 (R Core Team, 2016) with data filtering and Viterbi algorithms written in C++ using Rcpp (Eddelbuettel and François, 2011).

### 2.3 Model evaluation

The performance of the model in predicting daily time spent at the feeder was evaluated for each cow in the validation dataset using accuracy, sensitivity and specificity of the model predictions. We also calculated the number of daily visits and bout durations from the Insentec feeder visit raw data, HMM and raw logistic regression output.

## 3 Results

Cows were significantly more likely to have their head the in the feeding trough when located close to the feed barrier despite clear overlap in positioning measurements from feeding and not feeding cows (Figure 2a).

**Fig. 2.**
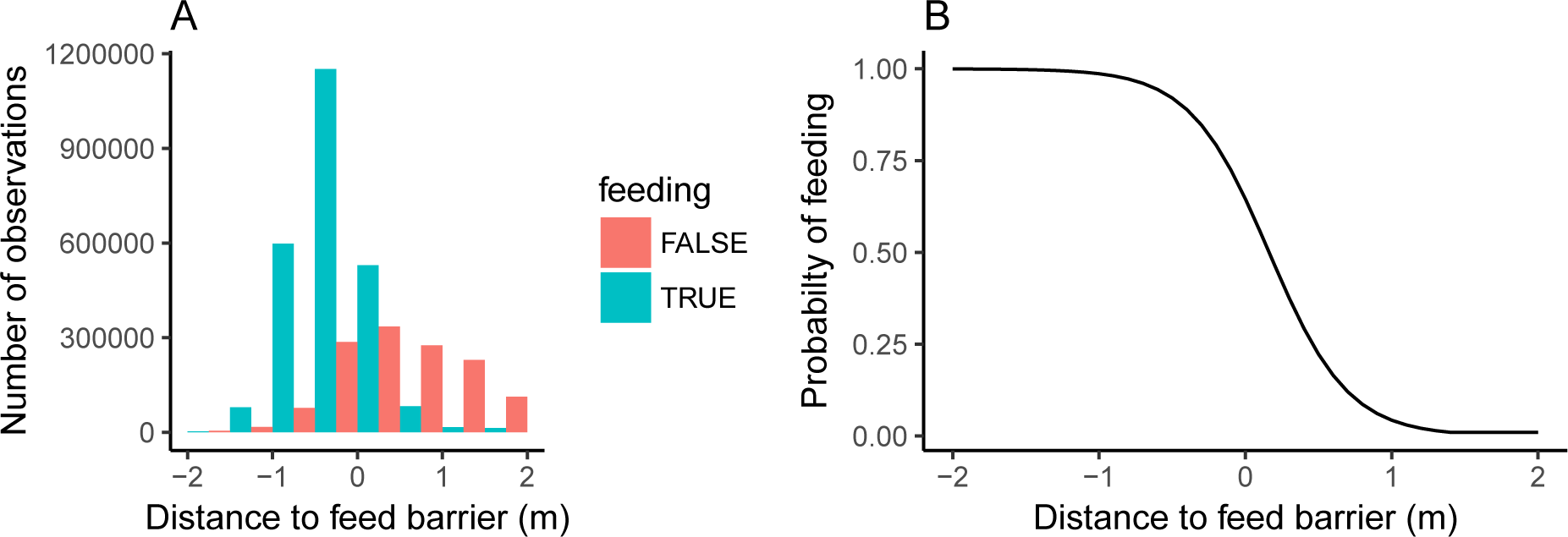
A) Histogram of observations from feeding and not feeding cows around feed barrier B) The probability of presence at the feeder as function of distance to feed barrier from the fitted logistic regression model

The probability of a cow being at the feeder obtained from logistic regression model fitted on the training data was:

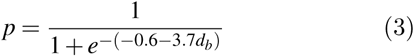

where:
p = probability of cow’s head being in feeding trough
d_b_ = distance to feed barrier in meters

The developed model was able to predict whether the cow was at the feeding trough or not with the accuracy of 97.6±0.01%, sensitivity 95.3±0.04% and specificity 97.9±0.01%. The figures are reported as the mean±standard deviation for cows in validation dataset.

The use of Viterbi algorithm clearly improved the model performance on measuring visit bouts compared to simple logistic regresson. Figure 3 shows feeding behavior of a cow measured using Insentec feeding throughs, predicted using logistic regression model and after using the viterbi algorithm to calculate the most likely state. The mean duration and the number of feeding bouts for validation dataset calculated from Insentec data and from both models is shown in Table 1.

**Fig. 3.**
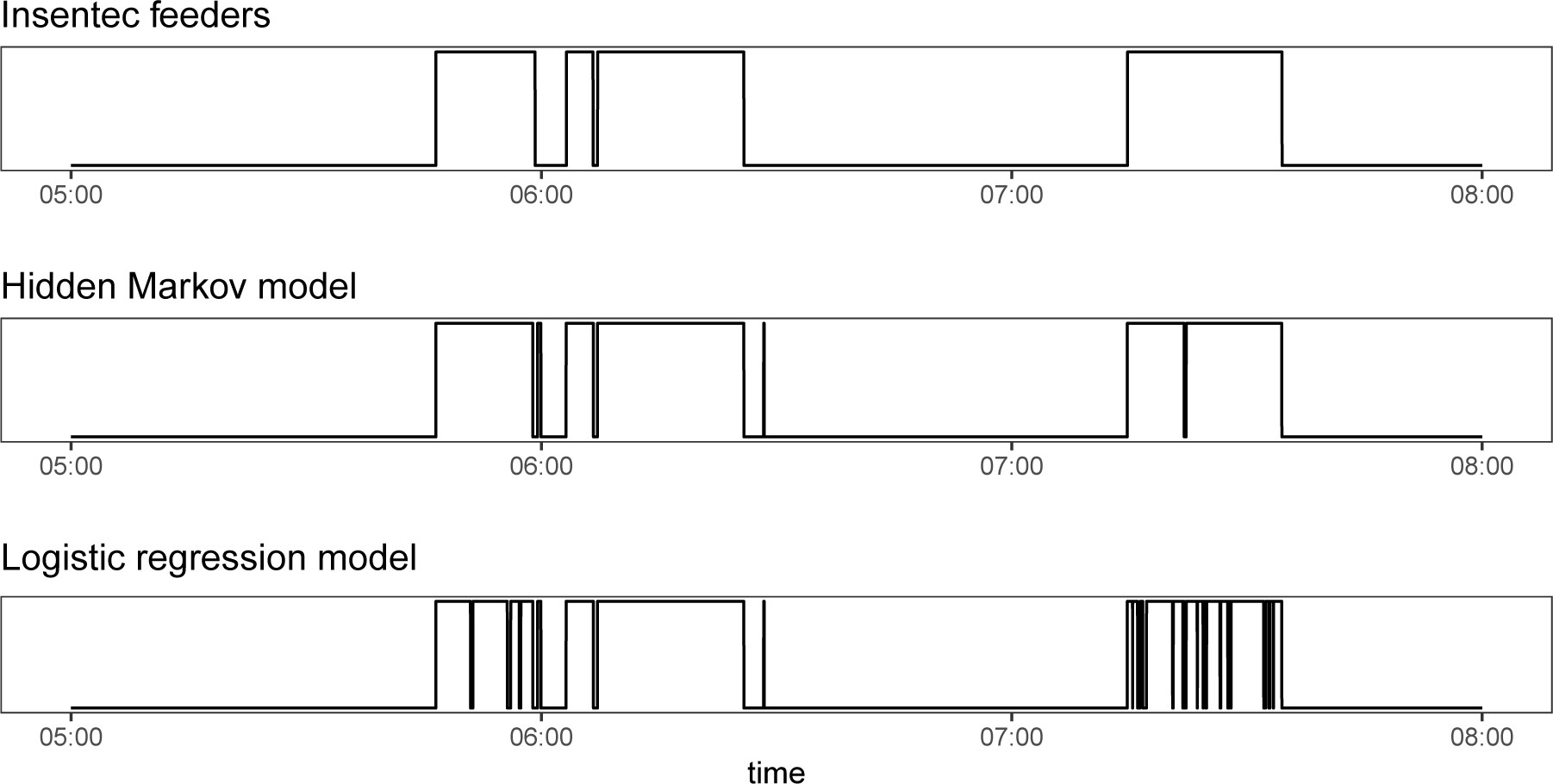
Measured (Insentec feeders) and predicted (Hidden Markov model and Logistic regression) presence at a feeding trough during 3 hours for a single cow. Top egde of figures represents feeding and bottom egde not feeding.

**Table 1.** Mean visit bout duration/cow and mean number of bouts/cow/day in validation dataset calculated from Insentec feeder data, using hidden Markov model (HMM) and logistic regression (LR)

| Method | Bout duration (s) | Bouts/day |
| --- | --- | --- |
| Insentec | 353 ± 123 | 31.8 ± 9.2 |
| HMM | 338 ± 101 | 34.5 ± 10.0 |
| LR | 286 ± 75 | 38.8 ± 10.8 |

## 4 Discussion and conclusions

We developed a model to the measure the feeding behavior of dairy cows using UWB indoor positioning system with 97.6% accuracy compared to reference method Insentec feeders (RFID and photocell combination). The results are in agreement with those obtained previously by Tullo et al. (2016) and Oberschätzl et al. (2015).

The use of Viterbi algorithm in addition to logistic regression clearly improved the model performance in calculating the duration and number of feeding bouts. The hidden Markov model still overestimated the number of bouts and underestimated the duration of feeding bouts on average, but the results were significantly closer to true values than raw predictions from logistic regression model.

The use of automated reference method allowed using a larger dataset than using the traditional method of behavioral observations and enabled the calculation of transition and emission probabilities with high accuracy. The same equations and fitted parameters can likely also be used to measure the presence at the feeder in other barns with good accuracy as long as the cow’s need to put their head trough the feeding barrier in order to have access to the feed. This obviously needs to be validated in any given environment before trusting the results.

A limitation of the method is that the time that the cow spends with the head in the feeding trough or area does not necessarily mean that the animal is actually eating. The time spent at the feeder without eating may very possibly change depending e.g. on the frequency of feed delivery and the feeding space allocated for each animal. However, time at the feeder itself has been shown to be a useful indicator of animals status (Norring et al., 2014; Urton et al., 2005).

UWB based positioning provides a good alternative for measuring time at the feeder and feeding bouts with comparable accuracy to RFID (our refence method) and computer vision based methods (Porto et al., 2015).

## References

Arcidiacono, C., Porto, S.M.C., Mancino, M., Cascone, G., 2017. Development of a threshold-based classifier for real-time recognition of cow feeding and standing behavioural activities from accelerometer data. Computers and Electronics in Agriculture 134, 124–134. doi:10.1016/j.compag.2017.01.021.

Eddelbuettel, D., François, R., 2011. Rcpp: Seamless R and C++ integration. Journal of Statistical Software 40, 1–18.

Halachmi, I., Meir, Y.B., Miron, J., Maltz, E., 2016. Feeding behavior improves prediction of dairy cow voluntary feed intake but cannot serve as the sole indicator. animal 10, 1501–1506. doi:10.1017/S1751731115001809.

Norring, M., Häggman, J., Simojoki, H., Tamminen, P., Winckler, C., Pastell, M., 2014. Short communication: Lameness impairs feeding behavior of dairy cows. Journal of Dairy Science 97, 4317–4321. doi:10.3168/jds.2013-7512.

Oberschatzl, R., Haidn, B., Peis, R., Kulpi, F., Völkl, C., 2015. Validation of automatically processed position data for evaluation of the behaviour of dairy cows. LANDTECHNIK – Agricultural Engineering; Vol 70, No 1 (2015).

Pastell, M., Frondelius, L., Jarvinen, M., Backman, J., 2018. Filtering methods to improve the accuracy of indoor positioning data for dairy cows. Biosystems Engineering 169, 22–31. doi:10.1016/j.biosystemseng.2018.01.008.

Porto, S.M.C., Arcidiacono, C., Anguzza, U., Cascone, G., 2015. The automatic detection of dairy cow feeding and standing behaviours in free-stall barns by a computer vision-based system. Biosystems Engineering 133, 46–55. doi:10.1016/j.biosystemseng.2015.02.012.

Porto, S.M.C., Arcidiacono, C., Giummarra, A., Anguzza, U., Cascone, G., 2014. Localisation and identification performances of a real-time location system based on ultra wide band technology for monitoring and tracking dairy cow behaviour in a semi-open free-stall barn. Computers and Electronics in Agriculture 108, 221–229. doi: 10.1016/j.compag.2014.08.001.

R Core Team, 2016. R: A Language and Environment for Statistical Computing. R Foundation for Statistical Computing. Vienna, Austria.

Ruuska, S., Hamalainen, W., Sairanen, A., Juutinen, E., Tuomisto, L., Järvinen, M., Mononen, J., 2014. Can stealing cows distort the results of feeding trials? An experiment for quantification and prevention of stealing feed by dairy cows from roughage intake control feeders. Applied Animal Behaviour Science 159, 1–8. doi: 10.1016/j.applanim.2014.08.001.

Shane, D.D., White, B.J., Larson, R.L., Amrine, D.E., Kramer, J.L., 2016. Probabilities of cattle participating in eating and drinking behavior when located at feeding and watering locations by a real time location system. Computers and Electronics in Agriculture 127, 460–466. doi:10.1016/j.compag.2016.07.005.

Thorup, V.M., Nielsen, B.L., Robert, P.E., Giger-Reverdin, S., Konka, J., Michie, C., Friggens, N.C., 2016. Lameness Affects Cow Feeding But Not Rumination Behavior as Characterized from Sensor Data. Frontiers in Veterinary Science 3. doi: 10.3389/fvets.2016.00037.

Tullo, E., Fontana, I., Gottardo, D., Sloth, K.H., Guarino, M., 2016. Technical note: Validation of a commercial system for the continuous and automated monitoring of dairy cow activity. Journal of Dairy Science 99, 7489–7494. doi: 10.3168/jds.2016-11014.

Urton, G., von Keyserlingk, M.A.G., Weary, D.M., 2005. Feeding Behavior Identifies Dairy Cows at Risk for Metritis. Journal of Dairy Science 88, 2843–2849. doi:10.3168/jds.S0022-0302(05)72965-9.

Viterbi, A., 1967. Error bounds for convolutional codesand an asymptotically optimum decoding algorithm. IEEE Transactions on Information Theory 13, 260–269. doi: 10.1109/TIT.1967.1054010.

Zucchini, W., MacDonald, I., Langrock, R., 2016. Hidden Markov Models for Time Series: An Introduction Using R, Second Edition. Chapman & Hall/CRC Monographs on Statistics & Applied Probability, CRC Press.

